# Selenium-enriched rapeseed extract synergizes with chemotherapy drug cisplatin in inhibiting proliferation and promoting apoptosis of colorectal cancer cells

**DOI:** 10.64898/2026.07.06.736755

**Authors:** Xiaofei Duan, Yiqing Lu, Hongyu Zhou, Zi Zhang, Zhanan Zhou, Miao Wang, Zhenna Chen, Xiaoling Dun, Yang Zhu, Hanzhong Wang, Lixi Jiang

## Abstract

Chemotherapy treatment of colorectal cancers (CRC) using cisplatin (CDDP) encounters problems of drug resistance by the cancer cells and cytotoxicity to normal cells, highlighting the urgent need for joint therapeutical strategies. Selenium-enriched rapeseed extracts exhibit anti-cancer effects but the bioactive components and mechanisms remain unclear. Here, we applied different solvents to fractionate the extracts from Selenium-enriched rapeseed and found that the water extract (WE) fraction significantly enhanced the cytotoxic effect of CDDP on cancer cells but no damage on normal cells. HPLC-ICP-MS analysis revealed that methylselenocysteine (MSC) and selenocystine (SeCys_2_) were the main selenium speciation in WE. Through cell biology and integrative multi-omics analysis, we found a synergistic anti-CRC cell effect when combining CDDP with MSC, sulforaphane (SFN), celastrol (Cel), Indole-3-carbinol (I3C), α-linolenic acid (ALA) or linoleic acid (LA). We propose that the CDDP-WE combination treatment holds the promise for improving curative efficacy for chemo-refractory CRC patients in the future.

## INTRODUCTION

Colorectal cancer (CRC) is the most common cancer in most of countries^1^. It is also the leading cause of death and ranks second only to lung cancer in people^2^. There are various methods for treating colorectal cancer, including traditional surgery, radiotherapy, chemotherapy, targeted therapy, and immunotherapy^3–5^. However, the cytotoxicity of chemotherapy drugs to normal cells somewhat limited use in actual clinical application^6^. Therefore, developing natural products with precise targeting of colorectal cells and the ability to form synergistic effects with chemotherapy drugs to enhance anti-tumor effects together with low cytotoxicity have good prospects in scientific value^7^.

Since the discovery of the activity of Platinum-based drugs, one of the first generation drugs called CDDP has been used in cancer treatment usually in conjunction with other ways such as surgery^8^, radiation^9^or immunotherapy^10^. Although CDDP can be effective for treatment of CRC its clinical application is constrained particularly by nephrotoxicity^11^. Natural products extracts have long been a unique source in combination with chemotherapeutics for the treatment of cancer^12^.Combination strategies are essential not only to enhance synergistic anticancer efficacy but also to alleviate cytotoxicity in healthy tissues through dose reduction^13^.

Many diet-based natural products have emerged as complementary and perspective source of novel anticancer adjuvant/drugs due to their overall safety and synergism with conventional chemotherapy in various cancers. Selenium-enriched rapeseed is a kind of vegetables with medicinal value, rich in various active ingredients such as selenium, polyphenols^14^, and flavonoids^15^. The glucosinolates in the sprouts can produce isothiocyanates with anti-cancer effects in the body^16^. In addition, selenium-enriched rapeseed also have a certain protective effect on improving male infertility^17^ and alleviating cadmium-induced reproductive toxicity^18^. Selenium-enriched rapeseed attracts people’s attention on account of its ability of selenium (Se) enrichment. Selenocystine (SeCys_2_), methylselenocysteine (MSC), selenomethionine (SeMet), selenite (SeIV), and selenate (SeVI) which are considered to be the main forms in rapeseed as revealed by Zhan et al^19^. However, the active anti-cancer ingredients of selenium-enriched rapeseed and their possible underlying impacts and mechanisms on CRC remain largely unclear.

In order to identify the active anticancer chemical ingredients of selenium-enriched rapeseed and uncover their anti-cancer mechanisms either as a monomer or in combination therapy in CRC treatment, we selected two common human colorectal cancer cell lines-SW620 and HCT116 cells as models that have been widely utilized in studies of CRC^20^. Firstly, we optimized the extraction parameters including extraction solvent, solvent concentration, solid-liquid ratio and extraction time. Next, we identified the most effective fraction that exerted a synergistic anticancer effect in CRC cells model. Third, the selenium speciation and content in different fractions were quantitatively analyzed by HPLC-ICP-MS. Finally, the monomers of selenium-enriched rapeseed were tested for their respective anti-cancer efficacy and cytotoxicity, and evaluated the efficacy when each of the monomers was combined with CDDP in treating colorectal cells. We found that MSC, sulforaphane (SFN) and celastrol (Cel) and Indole-3-carbinol (I3C), α-linolenic acid (ALA), linoleic acid (LA) displayed synergistic effect with CDDP on overcoming the chemoresistance in various patients. Herein, we not only demonstrate a promising therapeutic approach by combining CDDP and selenium-enriched rapeseed in treating CRC but also promotes the recognition of selenium-enriched rapeseed as a “dietary therapy” in people’s daily life.

## MATERIALS AND METHODS

### Chemicals and Reagents

Ethanol, petroleum ether, ethyl acetate, and n-butanol were purchased from China National Pharmaceutical Group Chemical Reagent Co., Ltd (Beijing, China). Sodium sulphite (Na_2_SeO_3_), sodium selenate (Na_2_SeO_4_), SeMet, SeCys_2_, and MSC were purchased from the China National Institute of Standard Material (Beijing, China). The Cell counting kit-8 (CCK-8) assay kit was obtained from Biosharp Co., Ltd. (Beijing, China). Annexin V-FITC/PI apoptosis kit and the cell cycle detection kit was purchased from Elabscience Biotechnology Co., Ltd. (Wuhan, China). Cisplatin, 2,2’-azino-bis-3ethylbenzthiazoline-6-sulphonate (ABTS), reactive oxygen species (ROS), L-lactate dehydrogenase (L-LDH), ALA, LA, Cel, SFN and I3C were provided by Beijing Solarbio Science and Technology Co., Ltd (Beijing, China). Trypsin-EDTA (0.05%) was obtained from Genview (FL, USA). Mitochondrial membrane potential assay kit with JC-1,5-ethynyl-2ʹ-deoxyuridine (EdU) cell proliferation kit, Radio-Immunoprecipitation Assay (RIPA) lysis buffer was obtained from Beyotime Biotechnology (Nanjing, China).The reduced glutathione (GSH) assay kit (microplate method) was purchased from Nanjing Jiancheng Bioengineering Institute (Nanjing, China).

### Plant Materials

The selenium-enriched rapeseed flowering stalks were collected from the Oil Crops Research Institute, Chinese Academy of Agricultural Sciences (Wuhan, China). The collected flowering stalks were ground into powder, freeze-dried with a lyophilizer (Labconco, USA) and then stored at 4℃.

### Optimization of Extraction Conditions

Single-factor experiments were conducted to confirm the optimal conditions of solvent concentration, solid-liquid ratio and extraction time based on the efficacy of selenium-enriched rapeseed flowering stalks alone and in combination with cisplatin on SW620 cells.

### Bioactivity-guided Fractionation

After determining the optimal extraction conditions, the crude extract (CE) was obtained. Then four fractions: petroleum ether extract (PEE), ethyl acetate extract (EAE), n-butanol extract (BE), and water extract (WE) were obtained by liquid-liquid partitioning^21^. Subsequently, PEE and EAE were dissolved in DMSO while other fractions directly dissolved in culture medium. The most active fraction was identified via the Cell Counting Kit-8 assay and the ZIP synergy score.

### (HPLC-) ICP-MS analysis

Total selenium content and different selenium species in different fractions were detected using (HPLC-) ICP-MS system (Agilent 7900, Agilent Technologies, USA) according to the method22. 50 mg of freeze-dried extracts and 5 mL of nitric acid were added into DigiBlock (LabTech, Beijing, China) in order to detect the total selenium content. At first, digested the mixture at 85°C for 3 h, and then evaporated at 120°C to nearly dryness. Then dissolved with 3 mL deionized water and introduced into ICP-MS. 20 mg extracts, 2 mL protease XIV (10 mg/mL) and 18 mL ultrapure water were added into per tube to analyze selenium species. After 40-min ultrasound treatment, the supernatant was obtained through centrifugation (12000 rpm×15 min) and introduced into HPLC-ICP-MS for the separation and identification of five kinds of selenium species.

### Cell Viability Assay

SW620, HCT116, CT26.WT, MCF-7, Hela, HepG2 and Huh-7 cells were purchased from Wuhan Procell Life Technology Co., Ltd. (Wuhan, China), NCM460 was purchased from American Type Culture Collection. Cells were cultured in a 37 °C, 5% carbon dioxide incubator.The cell viability of SW620 and HCT116 cells treated with different fractions for 24 h and was analyzed using the Cell Counting Kit-8 assay. The IC50 values were calculated using GraphPad Prism 9.0 software (San Diego, CA, USA). The ZIP synergy score was predicted using the online SynergyFinder software (https://synergyfinder.fimm.fi).

### Analysis of Cell Death and Cell Cycle

SW620 and HCT116 cells were subjected to 24 h treatments and then were collected. The cell death ratio and the cell cycle was detected by using the Annexin V-FITC/PI Apoptosis Detection Kit and cell cycle assay kit, respectively. Both are analyzed by flow cytometry. And data were analyzed with Flow Jo v10 software.

### Detection of ROS, GSH and L-LDH

SW620 and HCT116 cells were exposed to various treatments for 24 h and then collected. The flow cytometer was used to analyze the ROS level. GSH in SW620 and HCT116 cells was detected by GSH kits. For L-LDH detection, the cells were treated according to the L-LDH assay kit. And then the detection of GSH was conducted by measuring absorbance at 405 nm, while the L-LDH levels were detected as the absorbance at 450 nm by a microplate reader (Scientific Multiskan Sky, Thermo, America).

### Cell Proliferation Assay

The proliferation rate of SW620 and HCT116 cells was determined using an EdU assay kit (Beyotime Biotechnology, China). Treated cells were collected, and then stained with Alexa Fluor 488 Azide (30 min) and Hoechst 33342 (10 min), and detected by Keyence BZ-800E.Figure 4A,J,5G (cell morphology) and Figure 6A-D (mitochondrial morphology) are from the same instruments.

### Transmission Electron Microscopy

SW620 and HCT116 cells were treated with different combination for 24 h. After treatment, the cells were collected by centrifugation and washed twice with PBS, and then fixed overnight at 4°C in 2.5% glutaraldehyde, washed with PBS twice, and stained with 1% osmium tetroxide for 1 h under gentle rotation. Then rinsed with dd H_2_O for 3 times, fix with 2% uranyl acetate for 10 minutes once. Subsequently, the cell samples were dehydrated using 50%, 70%, and 90% ethanol solutions for 15 minutes each, and treated with 100% anhydrous ethanol once for 20 min. The cells were then incubated in a 1:1 mixture of embedder and acetone for two hours, ultrathin sectioning and imaged using the Talos L120C TEM (FEI, Thermo Fisher Scientific) equipped with the fast 4k × 4k Ceta™ 16M camera (BM-Ceta) and Thermo Scientific Maps at Center of Cryo-Electron Microscopy at Zhejiang University.

### TUNEL Assay

According to the instructions of TUNEL apoptosis detection reagent, cells were tested with 3 replicates per group. After washing the cells in each well with PBS, briefly, fix the cells with 4% paraformaldehyde for 30 minutes. and washed with PBS twice. Next, incubate the cells in 0.3% Triton X-100 for 5 minutes. Add 100 μL of TUNEL detection solution to each group and incubate at 37 °C in the dark for 60 minutes. After washing with PBS twice, stained with Hoechst solution for 10 minutes. And then washed with PBS once again, observed images under Keyence BZ-800E.

### Colony-Formation Assays

SW620 and HCT116 cells were plated in 6 cm dishes and the medium was replaced every 2 days. The different combination treated for 24h and then was continued until cell colonies were observed under a microscope. Ultimately, the cells were stained with 0.1% crystal violet.

### JC-1 Staining Assay

After the specified treatments, the mitochondrial membrane potential was assessed by using the Enhanced Mitochondrial Membrane Potential Assay Kit (Beyotime Biotechnology, C2006). The measurement was performed via Keyence BZ-800E in accordance with the manufacturer’s instructions.

### Molecular Docking and Dynamics Simulation

The core target (such as PTGS2) was screened by network pharmacology, and the stability of its binding with active ingredients (such as ALA) was verified by molecular dynamics simulation. The 2D structure of the active ingredients was obtained from the PubChem database and subsequently imported into Chem3D 23.1.1 software to construct and optimize the 3D structure in mol2 format. The crystal structure of protein targets was obtained by screening high-resolution structures from the RCSB PDB database^23^, and redundant groups such as water molecules and phosphate groups were removed using PyMOL 2.6 software. After preprocessing, they were saved as PDB format files^24^. Use AutoDock tool to perform hydrogenation, dehydration, charge allocation, and rotatable bond definition between proteins and ligands^25^, and conduct molecular docking simulations using AutoDock Vina 1.2.5 software^26^. Set the coordinates and size of the docking box based on the protein activity pocket, and screen the optimal binding conformation using the binding free energy score. Visualize the interaction between Discovery Studio 2019 and PyMOL 2.6 software, and draw 2D interaction diagrams and 3D combination mode diagrams^24^.

According to the molecular docking results, we further investigated the binding effect between active ingredients and targets using molecular dynamics simulation experiments. Molecular dynamics simulations of the ALA-PTGS2 complex were performed using Gromacs 2022 for 100 ns^27^. In this study, Sobtop_1.0 (dev3.1) was used to generate a topological file of the GAFF2 parameters^28^ based on the ligand structure, and the charge distribution of the ligand was carried out using the RESP method to ensure that the charge distribution conforms to the physical and chemical properties. For receptor proteins, the Amber99SB ILDN^29^ parameters are used. In the simulation process, the TIP3P water model was used for system solvation, and the minimum distance from protein atoms to the edge of the box was set to 1.0 nm for solvation to ensure sufficient solvation of the system. Electrostatic interactions were handled using the Particle Mesh Ewald (PME) method, and the LINCS was employed. The system underwent an energy optimization process: first, 3000 steps of steepest descent optimization were performed, followed by 2000 steps of conjugate gradient optimization. A 100-ns molecular dynamics simulation was conducted at a constant temperature of 310 K and pressure of 1 bar, with a coupling constant of 0.1 ps. During the simulation process, the gmx rmsd, gmx rmsf, gmx hbond, gmx gyre, gmx sasa, and gmx sham tools from Gromacs 2022 were used to calculate the root mean square deviation (RMSD), root mean square fluctuation (RMSF), hydrogen bond number, and cyclotron radius (Rg), respectively. Solvent accessible surface area (SASA) and free energy landscape map are used to analyze the stability, structural changes, and solvent effects of the system.

### LC-MS/MS Analysis

Cell samples were obtained after 24 hours of CDDP and WE alone or combination treatment.The tryptic peptides were dissolved in solvent A,The digested sample was loaded onto a home-made reversed-phase analytical column (25-cm length, 100 μm i.d.), following the protocol by the manufacturer.The peptides were subjected to capillary source followed by the timsTOF Pro mass spectrometry.The timsTOF Pro was operated in data independent parallel accumulation serial fragmentation (dia-PASEF) mode. The full MS scan was set as300-1500 (MS/MS scan range) and 20PASEF (MS/MS mode)-MS/MS scans were acquired per cycle. The MS/MS scan range was set as 400-850 and isolation window was set as 7 m/z.

### Determination of 11 Types of Fatty Acid Content

The extraction and determination method of fatty acids were improved from the previous study by Zhu et al.^30^. After gained freeze-dried powder, take about 500 mg and grind it, and then weigh about 50 mg of the powder. Add 2 mL of fatty acid extraction solution (chloroform: isopropanol, 2:1) and place it in the dark at room temperature for 2 hours. During this period, perform a 30 second vortex oscillation every half hour. After centrifugation for 5 minutes, take 500 mL of the supernatant and transfer it to another batch of labeled glass test tubes. Next add 2 mL of sulfuric acid methanol solution, vortex and centrifuge at 2500 rpm for 10 seconds, and then place it in an 80 °C water bath for 1 hour. After cool the sample and then add 2 mL of 0.9% NaCl solution and 1 mL of n-hexane, shake and centrifuge at 2500 rpm for 2 minutes. Next transfer the supernatant to a new glass test tube, then wash the residue twice with 2 mL of n-hexane, and merge the supernatant into the new test tube; After centrifugation (2500 rpm), take 700 μl and transfer it to injection bottle, either directly onto the machine or stored at -20 °C.The fatty acid content were measured by using GC-2014 (Shimadzu, Japan) gas chromatography.

### UPLC-MS/MS Analysis

UPLC-MS/MS was used place the extracts to identify the active ingredients of the WE.First of all, weigh grinding (30 Hz, 1.5 min) sample powder (30 mg) and add 1500 μL 70% methanolic aqueous internal standard extract. Then ultrasonic-assisted extraction was practiced. After centrifugation (rotation speed 12000 rpm, 3 min), the supernatant was gained and filtered through a microporous membrane (0.22 μm pore size) for subsequent UPLC-MS/MS analysis.

### Network Pharmacology

Firstly, the SMILE structural formulas of the major components of WE were retrieved in Pubchem (https://pubchem.ncbi.nlm.nih.gov/), and then use Swiss ADME tool (http://www.swissadme.ch) to predict the pharmacological properties of compounds and find potential core compounds. SwissTargetPrediction (http://www.swisstargetprediction.ch/) was employed to predict the targets of active compounds. Then using the keyword “colorectal cancer” in the GeneCards database (https://www.genecards.org/) and OMIM database (https://omim.org) to gain relevant targets, and continue analyzing the top 2000 targets. Next, remove duplicates. Using STRING database (https://string-db.org/)and resulting TSV-formatted data were analyzed by Cytoscape 3.9.1 to build a PPI database, and filter out core targets that are higher than the median. Then, a compound-target-function network was constructed to elucidate the intersection among active ingredients, core targets, and the function of anti-colorectal cancer. Preliminary screening of components with degree values greater than the median for further classification and analyze.

### RNA-seq Analysis

Total RNA of different treatment group was extracted by TRIzol (Invitrogen, CA, USA) and collected for RNA-seq analysis on the MGISEQ-2000 platform (MGI Tech Co., Ltd., Shenzhen, China). All experiments were performed in triplicate. Differentially expressed genes (DEGs) were obtained (p < 0.05) using DESeq2 (version 1.4). GO and KEGG were employed for the biological processes, cellular components, molecular functions, and pathway enrichment analysis.

### qRT-PCR Analysis

Total RNA was extracted from SW620 and HCT116 cells using Total RNA Isolation Kit (Vazyme Biotech, China). Complementary DNA (cDNA) were performed with All-in-one RT SuperMix (Applied Biological Materials, Canada). RT PCR was conducted by using ChamQ Universal SYBR qRT-PCR Master Mix (Vazyme Biotech) on a LightCycler 96 system (Roche, Basel, Switzerland). The primer sequences are provided in the supporting information.

### Statistical Analysis

Data was performed using GraphPad Prism 9.0 software (GraphPad Software, San 0Diego, CA, USA). Experiments were performed in triplicate and results are presented as mean ± standard deviation (SD). The differences between two groups were performed by t-test and one-way analysis of variance (ANOVA) followed between multiple comparison test. Statistical significance was p*< 0.05, p **< 0.01, and p ***< 0.001.

## RESULTS

### Outline of the experimental design

Figure 1 summarized the overall experimental flowchart described in this work. First, the fresh inflorescences of Selenium enriched rapeseed were harvested and ground into powders that were then placed in a freeze dryer for about 72 hours. After lyophilized, the dry powder was extracted with different concentrations of methanol or ethanol, which were called as crude extracts (CE). The CEs were checked for their effects on the different cancer cells alone or together with CDDP, respectively, to identify the most effective CE. The most effective CE was again lyophilized and the powder was dissolved in distilled water, and the yielded solution was then subjected to extraction sequentially with PEE, EAE and BE. The remaining solution after PEE, EAE and BE extraction was named as WE. The PEE, EAE and BE extraction fractions and remaining WE were checked for their effects on the cancer cells alone or together with CDDP, respectively.

**Figure 1.**
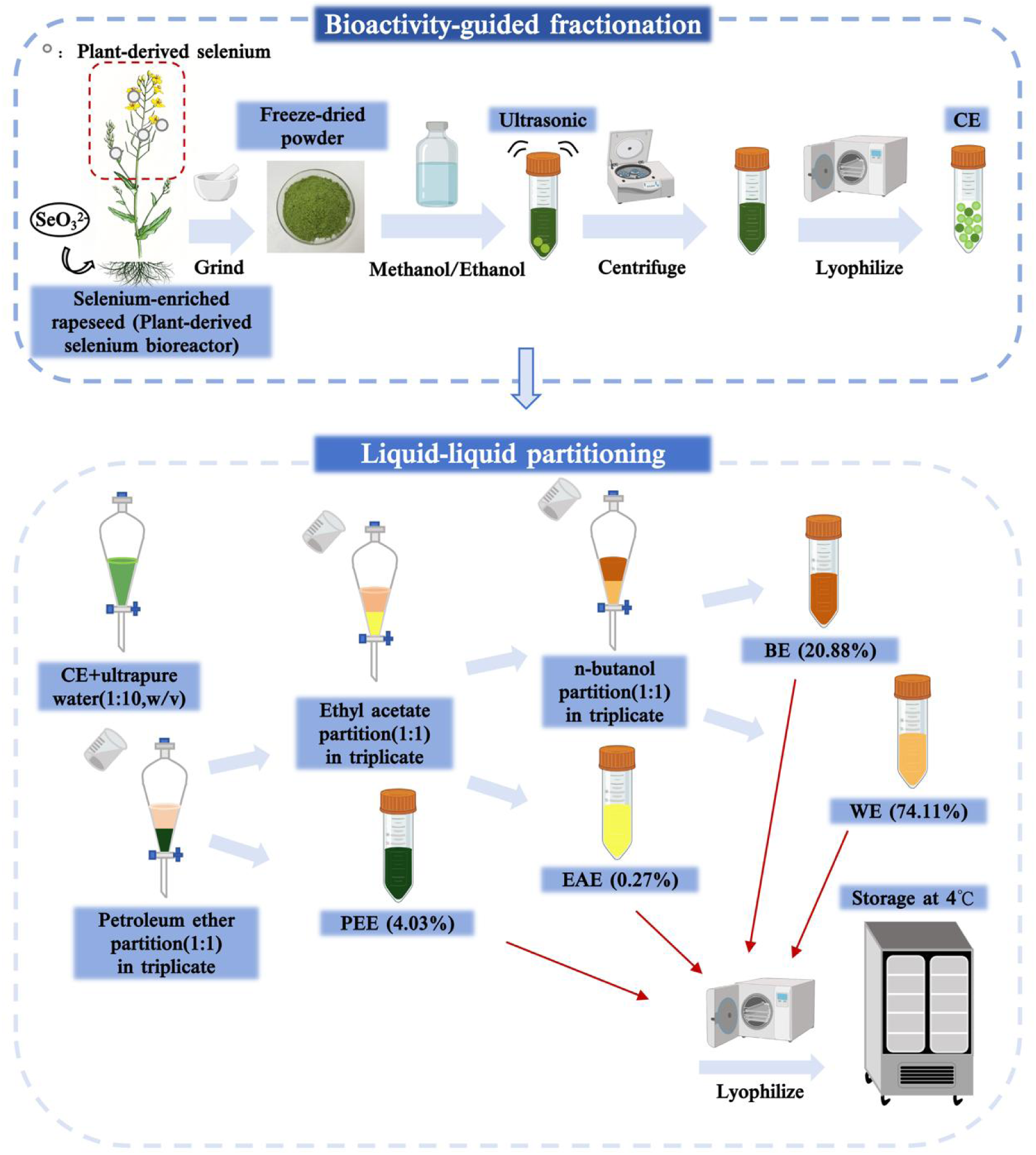
Diagram elucidates the flow chart of preparing crude extracts and subsequent fractioning using different solvents

### 50% Ethanol extracts were more effective in inhibiting cancer cell growth either alone or in combination with CDDP

To investigate the sensitivity of cancer cells to CDDP, an CCK-8 assay was conducted to assess the growth of various cell lines derived from human (MCF-7, HCT116, SW620, Hela, HepG2, and Huh-7) or mice (CT26.WT) following 24h treatment of CDDP at different concentrations. When compared with the control (without addition of CDDP), CDDP treatment obviously inhibited the growth of all tested cell lines in a dose-dependent manner, with IC50 values ranging from 0.77 μg/mL to 18.53 μg/mL (Figure 2A-B).

**Figure 2.**
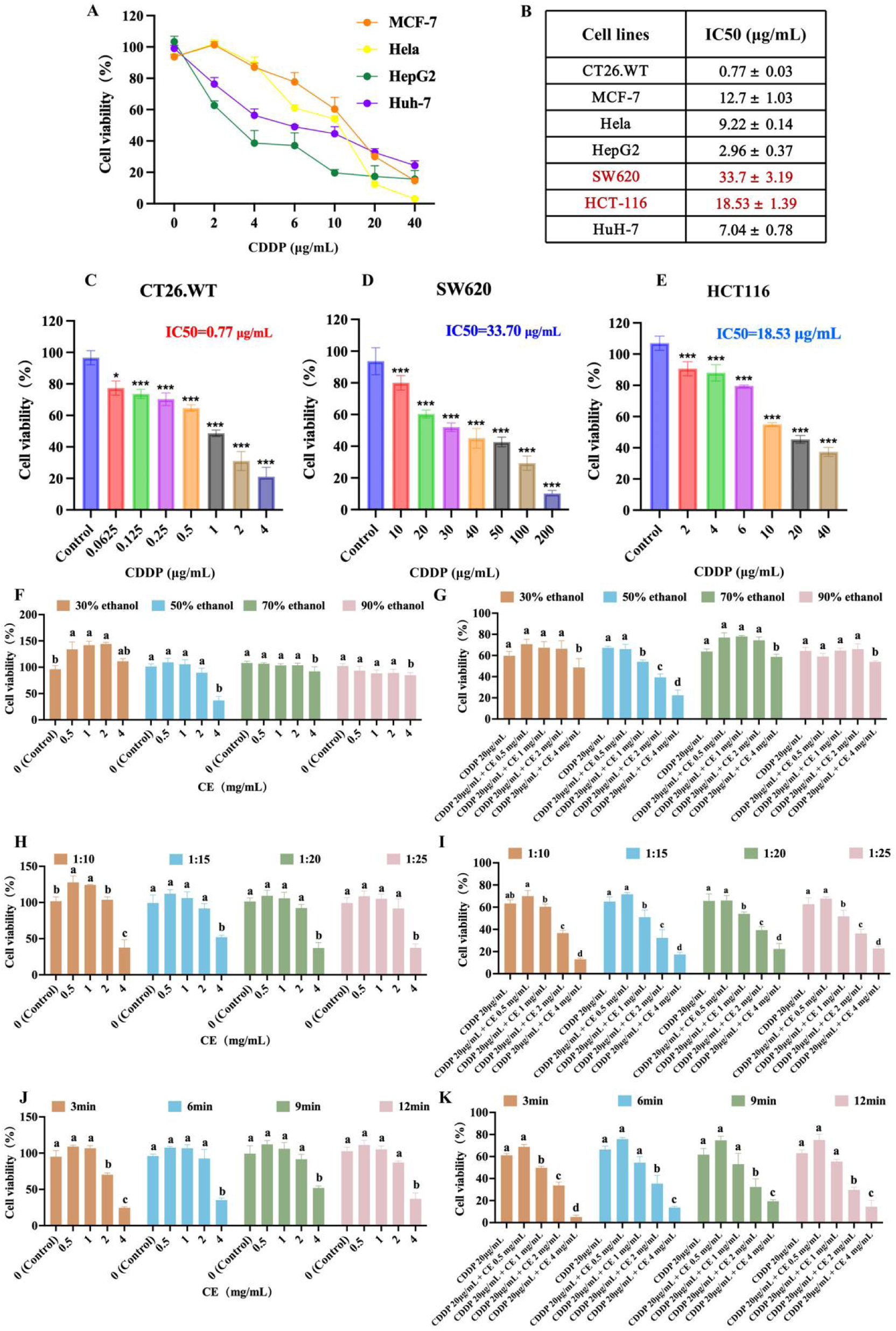
CE significantly enhanced the sensitivity of cancer cells to CDDP. (A) Cancer cell viability was evaluated using the CCK-8 assay after 24h treatment with different concentrations of CDDP. Control: without addition of CDDP. (B) IC50 values of CDDP in cancer cells as treated in (A,C). (C-E) CT26.WT, SW620 and HCT116 cell viabilities were assessed following CDDP treatment. The effects of ethanol concentration on the viability of cells treated with CE alone (F,G) and co-treated with CE and CDDP on the viability of SW620 cells. The effects of the solid–liquid ratio on the viability of cells treated with CE alone (H) and co-treated with CE and CDDP (I) on the viability of SW620 cells. The effects of the extraction time on the viability of cells treated with CE alone (J) and co-treated with CE and CDDP (K) on the viability of SW620 cells. The results are expressed as means ± SD (n = 3). For (F-K), * p < 0.05, ** p < 0.01, and *** p < 0.001 vs. control group. For (E,G,I,K), * p < 0.05, ** p < 0.01, and *** p < 0.001 vs. CDDP group. Different letters (a, b, c, d) indicate significant differences (p < 0.05)

We found that, in general, the human cancer cell lines exhibited stronger drug resistance compared to the mouse cancer cell line (Figure 2A,C-E). We also found that the CRC type cells SW620 and HCT116 exhibited relative stronger resistance to CDDP when compared with other kinds of cancer cells (Figure 2B). We therefore chose SW620 which showed the highest IC50 value as the cell model to determine which selenium-enriched rapeseed extraction could enhance the effect of CDDP. Since the half-maximal inhibitory concentration (IC50) of CDDP for SW620 cells was 33.66 μg/mL, we decided to use 20 μg/mL CDDP in the subsequent experiments.

Given that the choice of extraction solvent, solvent concentration, solid-liquid ratio and extraction time are crucial for ultrasonic extraction of bioactive compounds from nature products^21^, we assessed the impact of the extracts yielded from methanol and ethanol extraction with concentration ranging from 30% to 90% on cell viability either using the extract alone or in combination with CDDP. As shown in Figure 2F-G, no matter the solvent was methanol or ethanol, the extract alone showed inhibitory effects on SW620 cell proliferation at higher concentrations. Nevertheless, we observed that the 50% ethanol and 30% methanol extracts could enhance the effect of CDDP at lower concentrations. Because we noticed that the 30% methanol extract alone appeared to promote the growth of SW620 cells at low dosages. We thus chose 50% ethanol as the optimal extraction solvent for subsequent experiments.

As illustrated in Figure 2H and 2I, the 50% ethanol extract obtained from the solid-liquid ratio of 1:10 (the weight of the powder versus the liquid volume) promoted the cancer cell growth at the concentration 0.5 and 1.0 mg/ml. In contrast, the extracts alone from using the solid-liquid ratio ranging from 1:15 to 1:20 at the concentrations of 0.5, 1.0 and 2.0 mg/mL had almost no effect on the cell viability, but showed a significant growth inhibitory effect at 4 mg/ml. Importantly, we found that, when in combination with CDDP, the extracts at the concentrations of 2.0 and 4 mg/ml significantly enhanced the sensitivity of cancer cells to CDDP. Taken together, considering both cost and cell activity, we decided to take the 1:15 extracts as the optimum solid-to-liquid ratio for further experiments.

Next, the effect of extracts of different ultrasonic duration (3-12 min) on cell viability was studied. Figure 2J-K demonstrated that, with the duration of 3 min sonication, the extract concentration in the range of 2-4 mg/mL significantly enhanced the toxicity of CDDP towards SW620 cells. To save extraction time and ensure the effectiveness of combination on cell viability, 3 min sonication was chosen in the following experiments.

### WE Fraction Contains Bioactive Compounds for Inhibiting the Growth of the CRC-type Cells

The lyophilized 50% ethanol extracts were dissolved in the distilled water (1:10 w/v). We first assayed the effect of the water redissolved CE on the growth of seven types of cancer cells (CT26.WT, MCF-7, HCT116, SW620, Hela, HepG2, and Huh-7) and a human colon mucosal epithelial cell line NCM460 (Figure 3A). The IC50 for each type of cells are listed in Figure 3B. To further define the active ingredients, we used four different extraction fractions prepared according to Figure 1, namely PEE, EAE, BE and the remaining water redissolved CE (named as WE). We then assessed the possible synergistic effects of these fractions in combination with CDDP on the growth of SW620 and HCT116 cells by the CCK-8 assay. We quantified the synergistic effects by ZIP synergy scores. A ZIP Synergy score >0 indicated synergy, while a score >10 denoted strong synergy, as calculated by SynergyFinder software^22^ (Figure 3C). We observed that while all extraction fractions, when alone, showed inhibitory effects at higher concentrations, their combination with CDDP significantly enhanced the inhibitory effect on the proliferation of SW620 and HCT116 cells.To be specific, combinations such as CDDP-WE (ZIP: 25.32), CDDP-PEE (ZIP: 19.3), CDDP-EAE (ZIP: 10.02), CDDP-BE (ZIP: 35.67) in SW620 cells and CDDP-WE (ZIP: 10.81), CDDP-BE (ZIP: 22.5) in HCT116 cells exhibited an obvious synergistic inhibition, while CDDP-PEE (ZIP: -20.63) and CDDP-EAE (ZIP: -3.54) did not show a synergetic effect in HCT116 cells (Figure 3D-E).

**Figure 3.**
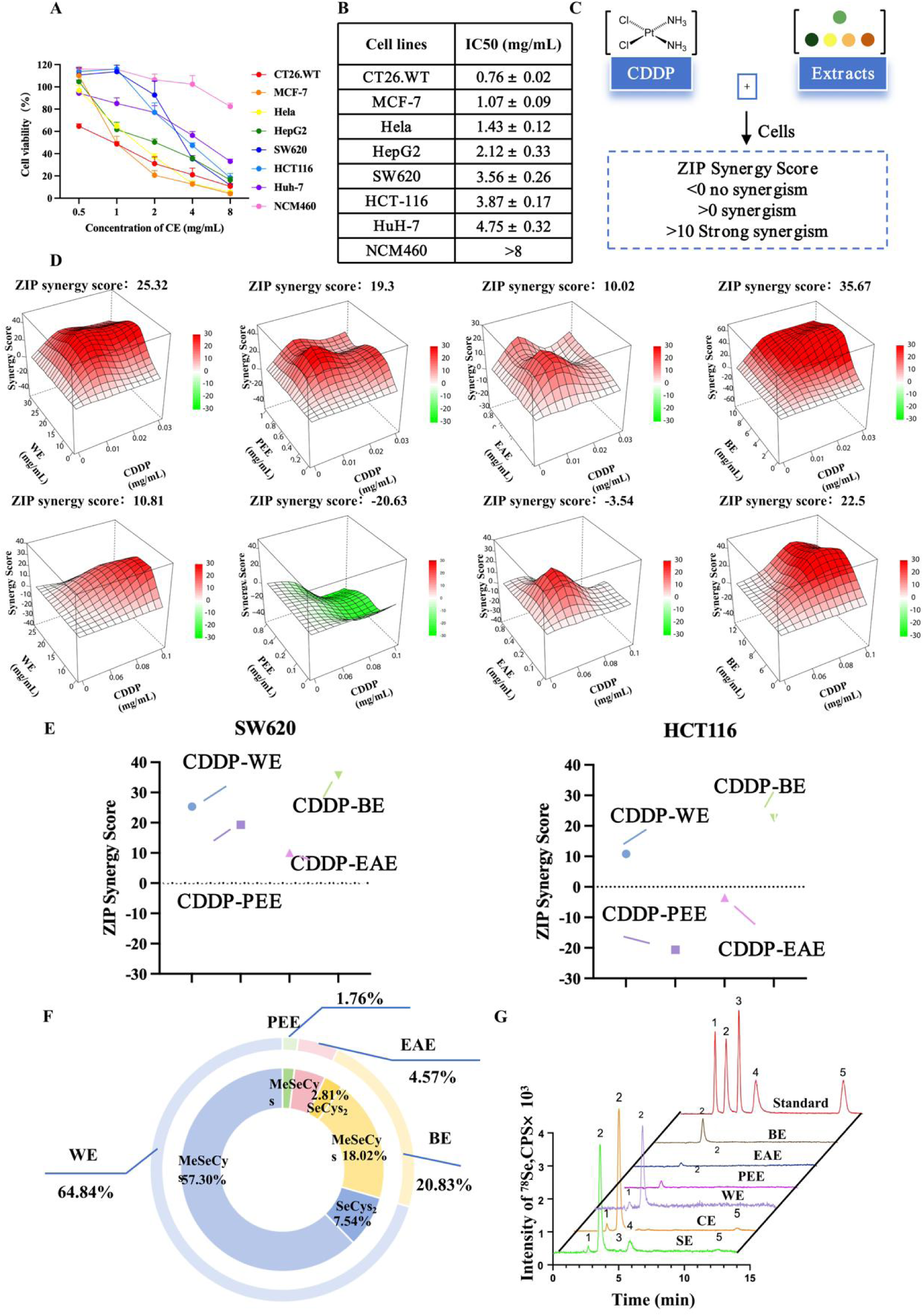
Evaluation of synergies between different extracts and CDDP on their abilities against cancer cells in vitro. (A) The CCK-8 assays assess the effect of the water redissolved lyophilized 50% ethanol CE alone on the proliferation of different types of cancer cells and the cytotoxicity of colon mucosal epithelial cell line NCM460 treated for 24 h at the indicated concentrations. (B) IC50 values of periplocin in CRC cells and NCM460 cells as treated in (A). (C) Study design. SW620 and HCT116 cells were treated with CDDP and WE for drug sensitivity. (D,E) The synergistic effects were quantified using the ZIP synergy scores (ZIP Synergy scores greater than 0 were considered indicative of synergism, and scores above 10 were considered strongly synergistic) via the SynergyFinder software, based on the cell viability from different extracts combinations. (F) Analysis of total selenium content in PEE, EAE, BE, and WE. (G) HPLC chromatograms of five selenium species in standard solutions, CE, PEE, EAE, BE, and WE. Peaks 1–5 in the chromatograms correspond to SeCys_2_, MSC, Se (IV), SeMet, and Se (VI), respectively.

We then analyzed total selenium contents in different fractions by subjecting the acid-digested fractions to the ICP-MS. As shown in Table S1, the WE fraction contained 168.61 ± 5.09 μg total selenium per gram of WE, accounting for 64.84 % of the total selenium in CE (Table S1). Next, we investigated the chemical speciation of selenium in the fractions (CE, PEE, EAE, BE, and WE). The results of HPLC-ICP-MS showed that five selenium species were in the fractions (Figure 3F-G). Interestingly, neither SeCys nor Se (VI) was detected in any fraction. SeMet is only detected in CE. These results showed that MSC was the predominant selenium species in CE and was largely retained in WE (74.11 %). 11 fatty acids of SE were identified by GC (Table S2). Taking into account of the selenium compositions and the effect on SW620 and HCT116 cells, WE was selected for the subsequent experiments. The CDDP-WE combination displayed significantly synergistic effect compared to their single use. In summary, the CDDP-WE combination is considered as a combinatorial strategy with anti-colorectal cancer potential.

### WE Enhances the CDDP-induced Oxidative Stress in CRC Cells

To assess the effects of the CDDP-WE combination on SW620 and HCT116 cells, we observed cell morphological changes caused by different treatments (Figure 4A). The CDDP-WE combination treatment reduced cell proliferation and increased the proportion of cell debris, suggesting a combinatory antiproliferative and cytotoxic effect. In order to further investigate the mechanism of action of the CDDP-WE combination on CRC cells, we measured the ROS levels after the CDDP-WE treatment. Compared with WE treated alone, the ROS levels in SW620 and HCT116 cells were elevated significantly after the CDDP-WE combination treatment (Figure 4B-E). These results suggested that the CDDP-WE combination inhibits CRC cell growth by promoting the ROS accumulation. Lactate dehydrogenase (LDH) release assay revealed a notable increase in the LDH levels in the CDDP-WE combination group, indicating cell membrane rupture and CRC cell death (Figure 4F,H). The addition of WE led to depletion of GSH in cancer cells, exacerbated oxidative stress damage induced by CDDP, and improved the therapeutic effect of CDDP (Figure 4G,I). Moreover, our results also indicated the possibility of using the EC50 for ABTS to evaluate the extracellular antioxidant activity of WE and showed the good free-radical-scavenging capabilities of WE^23^.

### The CDDP-WE Combination Treatment Inhibits CRC Cell Proliferation and Promotes Cell Apoptosis

The EdU labeling assay revealed that the CDDP-WE combination exhibited a significant synergistic effect on inhibiting cell proliferation (Figure 4J). Treating SW620 and HCT116 cells with the CDDP-WE combination shifted the distribution of cells from the G0/G1 phase to the S and G2/M phase as revealed by flow cytometry analysis. The proportion of S phase cells in the CDDP-WE treatment groups increased obviously whereas the WE alone treatment did not induce any changes in the cell cycle phases (Figure 5A-B). The results indicate that WE sensitized the CRC cells to CDDP through cell cycle arrest in the S phase, thereby inhibiting cell proliferation. To further evaluate the antiproliferative effects, colony formation assays were performed. The CDDP-WE treatment significantly reduced colony formation compared with single treatments and this effect was significantly better than did the CDDP alone treatment (Figure 5C-D).

**Figure 4.**
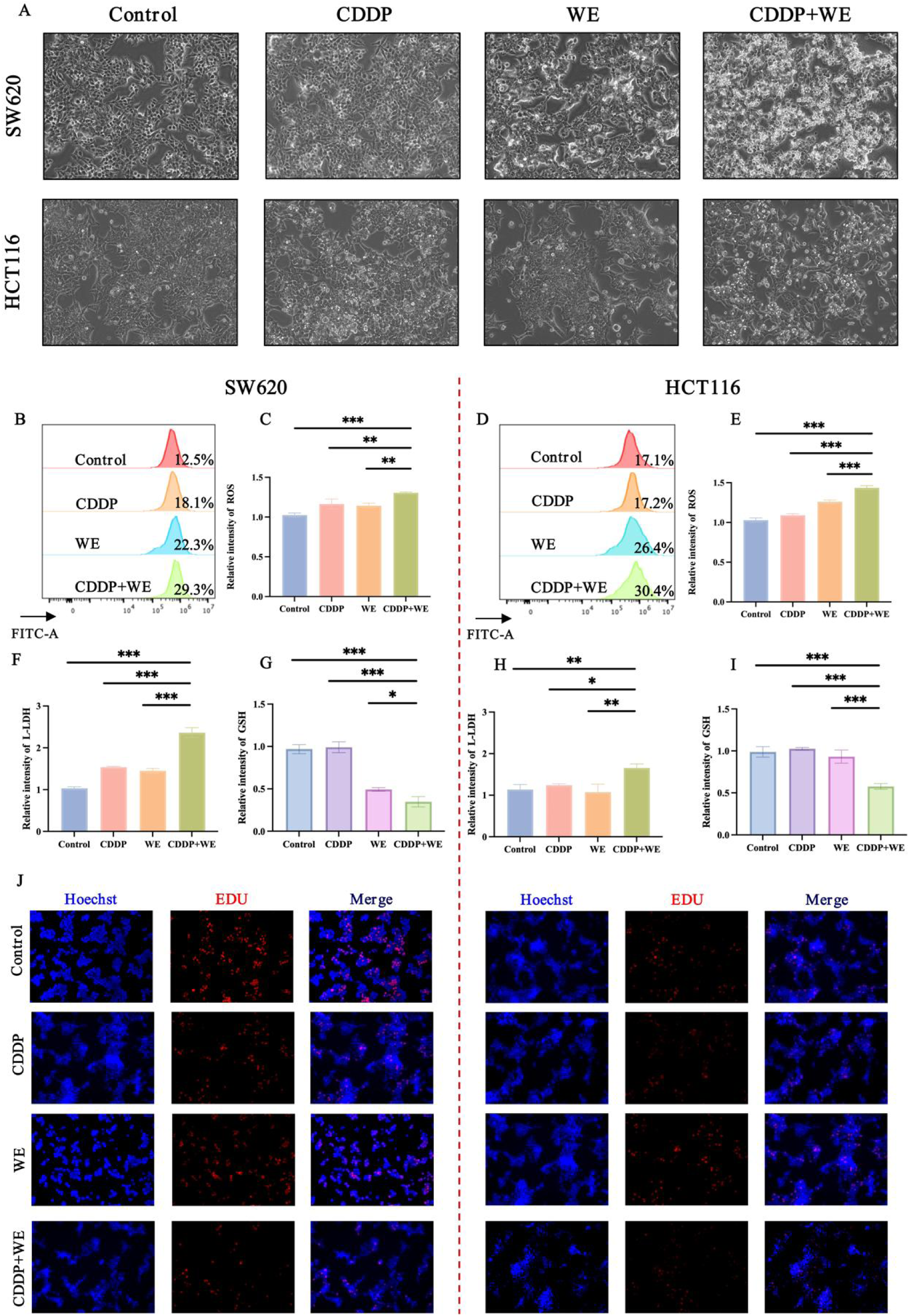
The CDDP-WE combination treatment synergistically improves antioxidant capacity and induces cell death. (A) Cell morphological of the SW620 and HCT116 cells after the CDDP-WE combination treatment for 24 h. (B-E) Measurement of the ROS release. (F,H) Measurement of the LDH release. (G,I) Measurement of the GSH release. (J) Fluorescent staining for EdU assay. The mean ± SD is shown, n = 3. * p < 0.05, ** p < 0.01, *** p < 0.001.

**Figure 5.**
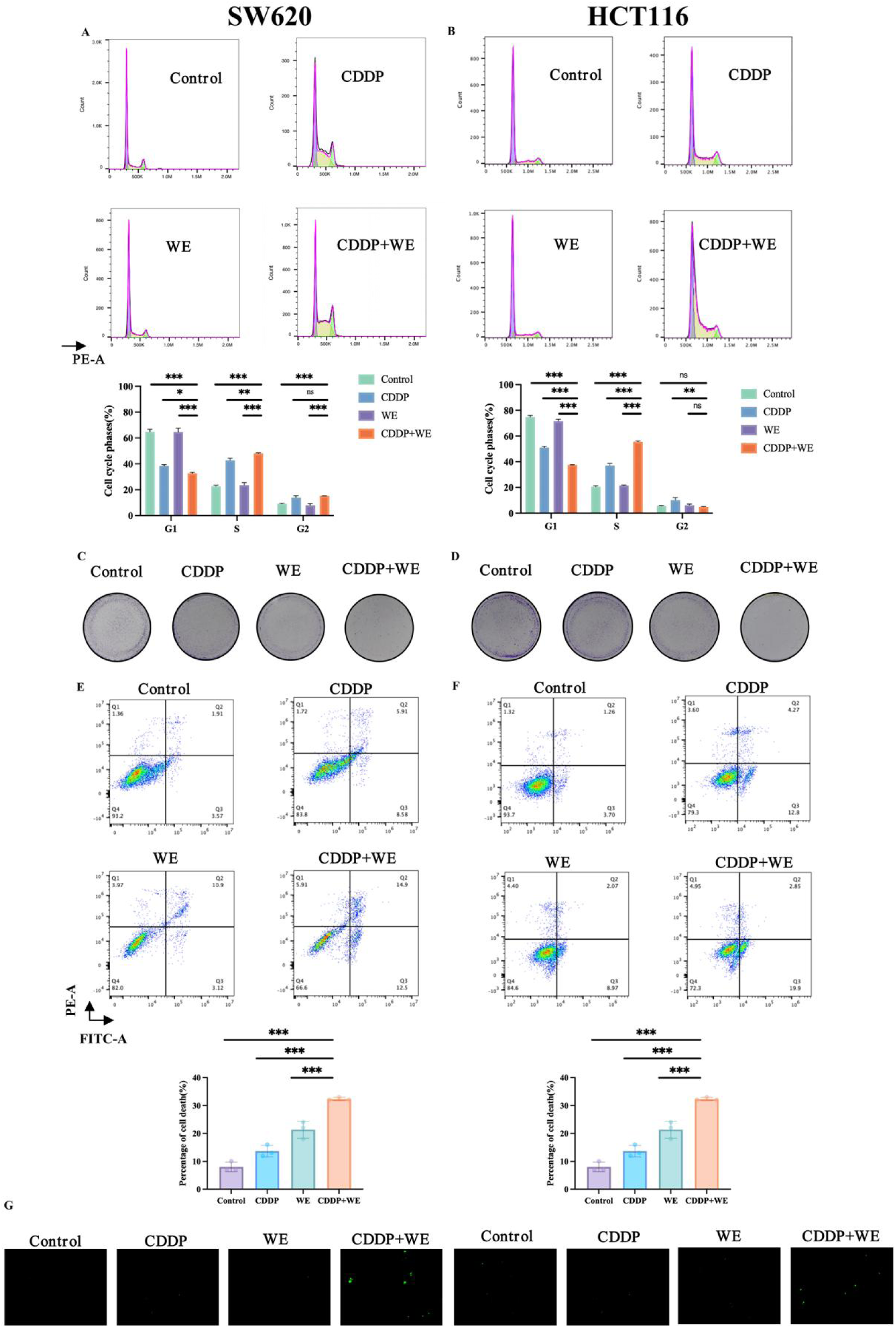
The CDDP and WE combination treatment synergistically inhibits cell proliferation and triggers apoptosis in CRC cells. (A-B) CDDP-WE influenced the cell cycle in HCT116 and SW620 cells, the cell cycle was analyzed by PI staining and detected using flow cytometry. (C-D) Colony-forming after the CDDP and WE combination treatment for 2 weeks in HCT116 and SW620 cells. Colony formation was fixed in methanol, stained with crystal violet. (E-F) Cell death rate was assessed by flow cytometry after staining with phartmingen Annexin V-FITC Apoptosis Detection Kit in HCT116 and SW620 cells following CDDP and WE treatment for 24 h. (G) Fluorescent images obtained using a fluorescence microscope. Apoptosis-induced cells released green fluorescence upon analysis with the DeadEnd™ Fluorometric TUNEL System. The mean ± SD is presented, n = 3. * p < 0.05, ** p < 0.01, and *** p < 0.001 vs. different group.

Next, we attempted to investigate the anti-cancer function of the CDDP-WE combination by analyzing cell viability. We adopted the flow cytometry to analyze cell death through Annexin V/PI double staining. The percentage of apoptotic SW620 and HCT116 cells significantly increased after incubation with the CDDP-WE combination, reaching 34% and 28%, respectively (Figure 5E-F).

The TUNEL staining results showed that there were very few apoptotic cells in the control group, while the fluorescence intensity of apoptosis was significantly increased in CDDP-WE groups. A large number of green apoptotic cells were observed in the fluorescence images (Figure 5G), suggesting that, in addition to arrest cell proliferation, the CDDP-WE combination also effectively promotes apoptosis.

Taken together, the roles of the CDDP-WE combination in promoting ROS production, inhibiting cell proliferation and triggering cell apoptosis are likely the main contributors towards its anti-CRC effect.

### The CDDP-WE Treatment Synergistically Induces Apoptosis Through Mitochondrial-Dependent Pathway

To investigate whether mitochondrial dysfunction contributed to CDDP-WE-induced apoptosis, the SW620 and HCT116 cells were analyzed by transmission electron microscopy (TEM). TEM showed that, compared with the control groups, the CDDP-WE combination treated SW620 and HCT116 cells exhibited a deformed mitochondrial morphology such as swollen mitochondria cristae ambiguity and tumidness (Figure 6A,C). An increasing number of studies indicate that the breakdown of MMP (Mitochondrial membrane potential) may lead to mitochondrial dysfunction, which plays a dual role in natural products induced cell apoptosis^24^. In order to investigate whether the CDDP-WE combination treatment induced cell apoptosis occured through the mitochondrial apoptosis pathway, the JC-1 kit was used to detect changes in MMP on SW620 and HCT116 cells. Green fluorescence marked cells that had undergone apoptosis or necrosis, while red fluorescence marked alive cells that maintained normal mitochondrial membrane potential. As shown in Figure 6B,D, compared with the control group, the red fluorescence was also significantly reduced and the green fluorescence was enhanced, indicating a decrease in mitochondrial membrane potential coupled with an increase in apoptotic cells in the CDDP-WE group. Therefore, it appears that the CDDP-WE combination synergistically activates the apoptosis through the intrinsic mitochondrial apoptotic pathway in CRC cells.

**Figure 6.**
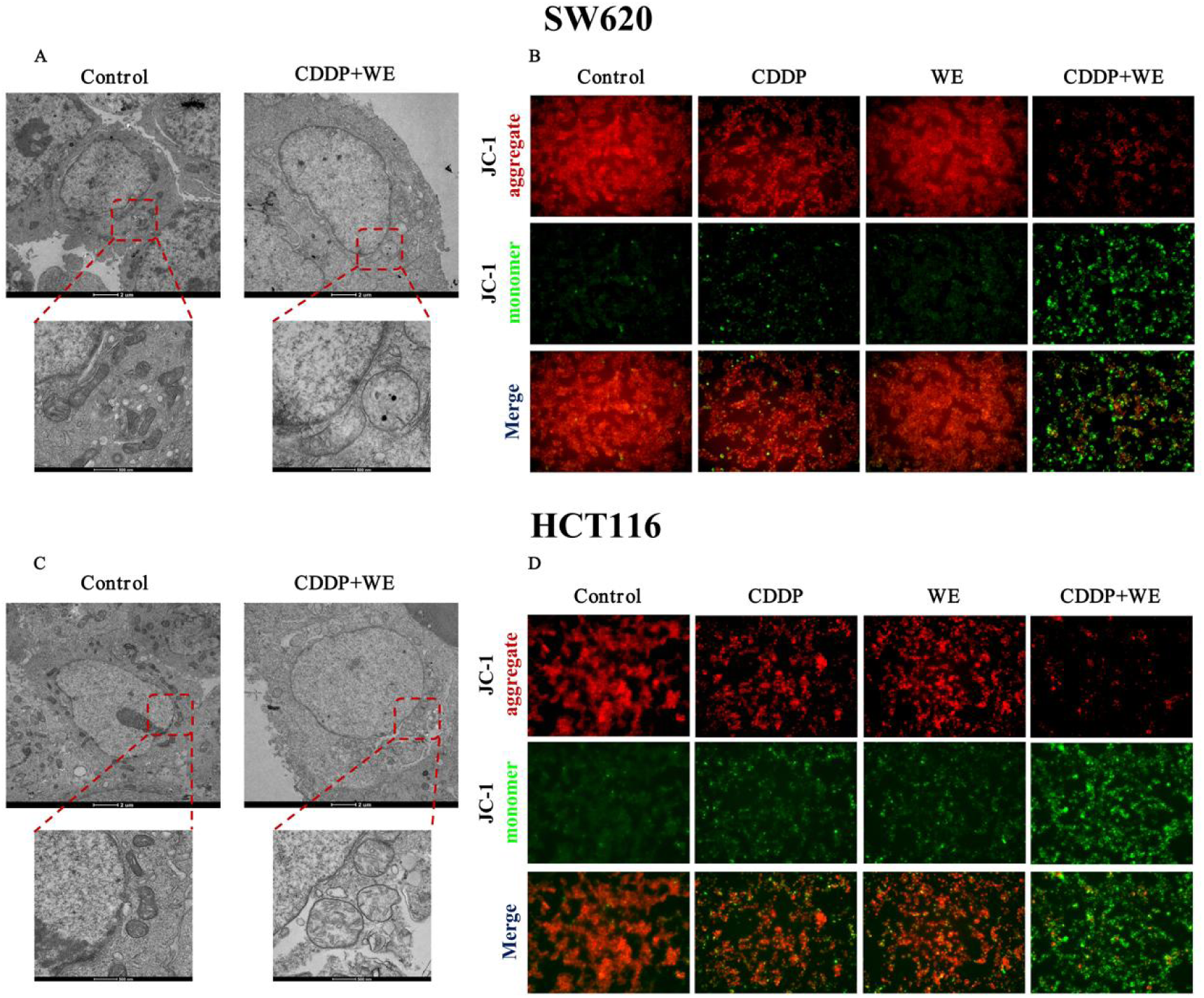
The CDDP-WE combination treatment activates the mitochondrial intrinsic apoptotic pathway. (A,C) TEM observation of the mitochondrial morphology. (B,D) Membrane potential detection of CRC cells.

### Characterization of the Chemical Profile of Selenium-Enriched Rapeseed

Next, we conducted a comprehensive analysis of the chemical composition of WE by employing the ultra-performance liquid chromatography-tandem mass spectrometry (UPLC-MS/MS). Selenium-enriched rapeseed is a classical source of natural products for identifying active ingredients with anti-cancer effects. Molecular formulas and chemical profile of WE were further analyzed through reference databases and tandem mass spectrometry interpretation. Through prediction, a total of 314 high-content chemical compounds in WE were identified, including classes:76 amino acids and derivatives, 46 benzene and substituted derivatives, 36 organic acids, 32 heterocyclic compounds, 29 others, 16 alcohol and amines, 16 alkaloids, 13 lipids, 12 phenolic acids, 11 flavonoids, 7 terpenoids, 6 FA, 4 GP, 4 nucleotides and derivatives, 4 lignans and coumarins, 1 quinones and 1 steroids (Figure 7A). The metabolome demonstrates the diversity and complexity of WE, so it is necessary to explore the main active ingredients in WE that synergistically exert anti-CRC effects with chemotherapeutics.

**Figure 7.**
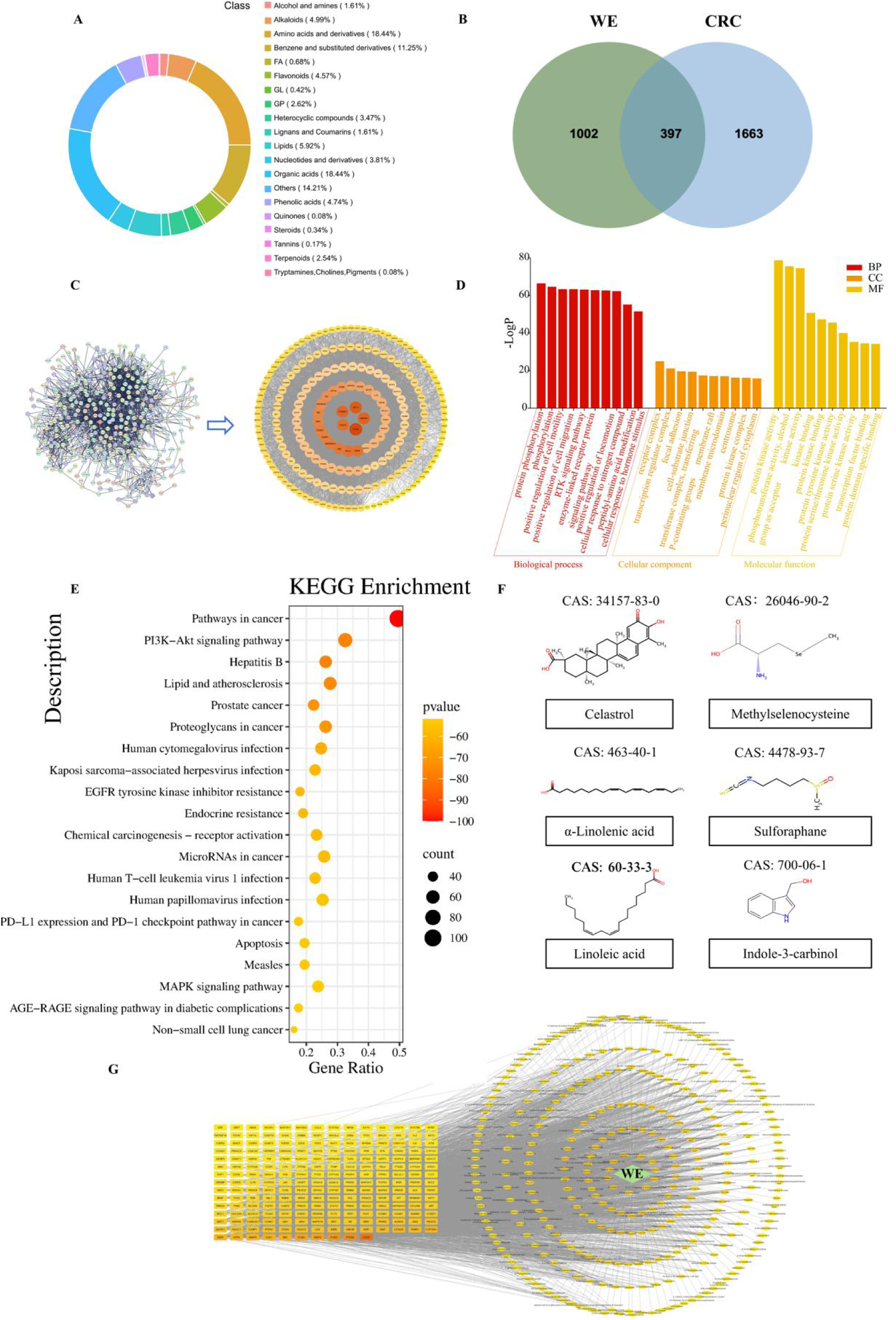
Network pharmacology analysis of WE in the treatment of colorectal cancer. (A) Structural classification of compounds contained in the WE. (B) Venn diagram of gene sets between groups. (C) PPI network. (D-E) GO and KEGG pathway enrichment analyses of DEGs. (F) Chemical structures of the compounds that considered as the basic active ingredients with anti-colorectal cancer potential. (G) Network of drug active components-intersection targets.

### Network Pharmacology Analysis of The Main Active Ingredients of WE in The Treatment of CRC

Network pharmacology has now become a key approach to exploring core targets of natural products. 327 compounds including 2 selenium-containing compounds identified via HPLC-ICP-MS, 314 non-selenium compounds determined by UPLC-MS/MS and 11 fatty acids detected by GC exhibiting bioavailability and drug-likeness were imported into the Swisstargetprediction database to predict their potential targets. A total of 1,399 compound-target interactions were identified. Using the keyword ’colorectal cancer’ in GeneCards data to retrieve relevant targets allowed to obtain 125,003 targets. After sorting them by score value and selecting the top 2,000 targets for further analysis, we obtained 96 targets in OMIM database. Consequently, we analyzed the intersection of targets and obtained 2,060 targets after deduplication (Figure 7B). With the confidence level of high confidence (0.900) and after deleting free node and sanding, we obtained 397 intersecting targets and these targets were entered into the STRING-database. Then, the resulting protein-protein interaction (PPI) network was introduced. And further analyzed and visualized using the CentiScaPe plug-in within Cytoscape, three centrality-measures (degree, closeness, and betweenness centrality) were calculated for each target. Then, the intersection was taken to obtain 206 core targets (whose values exceeded the respective medians) (Figure 7C). In this way, the targets identified included CDK2, PTGS2, ITGB3, Mitochondrial membrane potential 3, ITGB1, SRC, CDK1, MMP9, DPP4 and EGFR which were emerged as the top ten core targets, and these nodes are the representative of the WE key targets against CRC cells. In addition, the above 206 core targets were imported into the David database for GO and KEGG analysis, the pathways enriched by KEGG, included PI3K-Akt signaling pathway, apoptosis signaling pathway, MAPK signaling pathway, and pathways in cancer cells which are associated with cell apoptosis. GO enrichment analysis showed that the biological processes of WE in treating colorectal cancer mainly enriched on the regulation of protein phosphorylation and positive regulation of cell motility and migration. The cellular components mainly included receptor complex and transcription regulator complex; the molecular functions mainly focused on kinase activity, protein kinase binding, and transcription factor binding (Figure 7D-E). Through literature searching, it was found that Cel, MSC, ALA, SFN, LA and I3C have been reported to be used as synergistic chemotherapy drugs that enhance anti-CRC efficacy. Therefore, they were selected as the active ingredient (ligand) for molecular docking (Figure 7F). To elucidate correlations between functional active ingredients in WE and core targets linked to CRC, a compound-target-function network was constructed (Figure 7G).

BAX, BCL2, CASP3, CASP9, CDK2, HMOX1, JUN, MAPK1, MYC, STAT3, TP53, AKT1, CCNB1, CCND1, EGFR, FOS, GPX4, PTGS2, MMP3 appeared more frequently in the KEGG enriched pathway and are known to be associated with cell growth and development (Figure 8A). In addition, some of them are related to cellular oxidative stress. Therefore, these nineteen proteins were selected as protein targets for docking of the ligands.

**Figure 8.**
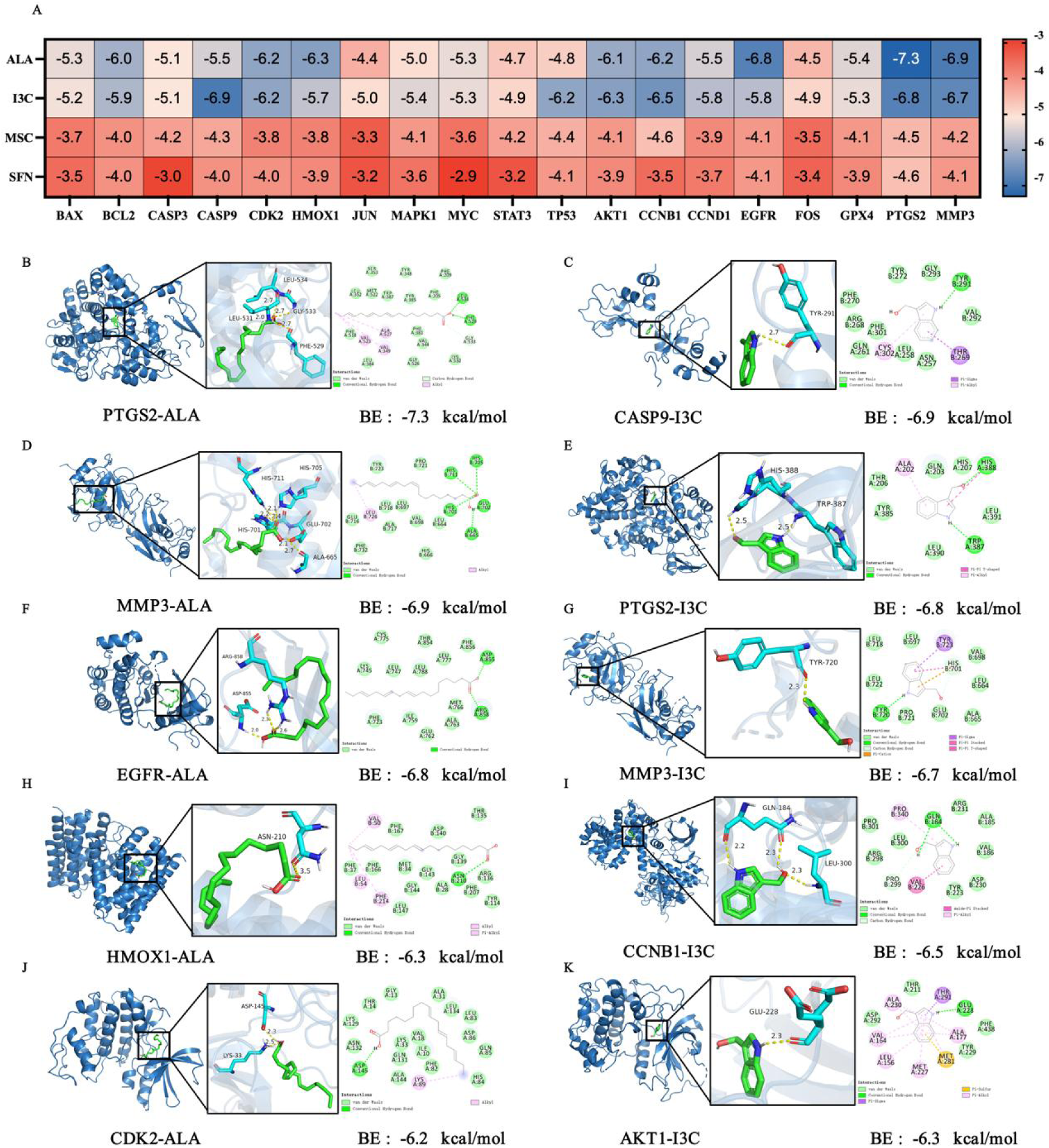
Molecular docking simulation of ALA and I3C interactions with core colorectal cancer-associated targets. (A) Binding affinity heatmap of ALA and I3C against 19 core targets, color-scaled by docking energy (kcal/mol). (B) ALA-PTGS2 protein molecular docking. (C) I3C-CASP9 protein molecular docking. (D) ALA-MMP3 protein molecular docking. (E) I3C-PTGS2 protein molecular docking. (F) ALA-EGFR protein molecular docking. (G) I3C-MMP3 protein molecular docking. (H) ALA-HMOX1 protein molecular docking. (I) I3C-CCNB1 protein molecular docking. (J) ALA-CDK2 protein molecular docking. (K) I3C-AKT1 protein molecular docking.

### Molecular Docking and Molecular Dynamics Simulations Reveal Stable Interactions Between Active Ingredients and Core Therapeutic Targets

Docking energy value of < -4.25 kcal/mol (1 cal ≈ 4.186 J) is generally acceptable for indicating binding activity between the molecules, whereas, <-5.0 kcal/mol and <-7.0 kcal/mol indicates good and strong binding activity, respectively. Based on network pharmacology enrichment results, we conducted molecular docking analysis of ligand-target to evaluate their affinities (Figure 8A). Through molecular docking thermography and visualization display, it was shown that ALA good binding affinity with fifteen core target proteins except JUN, STAT3, TP53 and FOS, while I3C has good binding affinity with seventeen core target proteins except STAT3 and FOS (Figure 8B-K). Among the components, ALA exhibited the strongest binding ability with PTGS2. The binding score between ALA and PTGS2 was below −7 kcal mol ^−1^. The LEU534 and PHE529 residues on the PTGS2 receptor form hydrogen bonding interactions with ALA, while the ALA527, VAL523, and VAL349 residues on the PTGS2 receptor form hydrophobic interactions with ALA. SER353, TYR348, PHE209, PHE205, TYR385, TRP387, MET522, LEU352, PHE518, LEU384, GLY526, PHE381, VAL344, GLY526, LYS532 and LYS532 residue forms a vander Waals interaction with ALA.

### The CDDP-WE Combination Treatment Activates Tumor Suppressive Pathways in the CRC SW620 and HCT116 Cells

To further investigate the molecular mechanisms underlying the CDDP-WE combination treatment induced CRC cell death, transcriptome analysis was performed. Comparative transcriptomics identified 5,301 downregulated and 5,604 upregulated genes in Control versus CDDP-WE groups in SW620, with Control versus CDDP-WE groups 5,328 downregulated and 5,184 upregulated transcripts in HCT116(fold change >0, p < 0.05) (Figure 9A,E). Cross-comparison revealed that 463 genes were upregulated and 739 downregulated in both SW620 and HCT116 cells (fold change >1, p < 0.05) (Figure 9B,F). Subsequent GO functional enrichment analysis (Figure 9C-D) and KEGG pathway enrichment analysis (Figure 9G-H) of the DEGs showed that pathways including apoptosis, P53 signaling pathway, MAPK signaling pathway, mTOR signaling pathway, AMPK signaling pathway and mitophagy were enriched in both SW620 and HCT116 cells, suggesting these pathways might serve as key pathways of the anti-CRC effect exerted by the combination therapy. The expression of key genes (*CASP6*, *CASP9*, *MAPK14*, *BAX* and *JUN*) related to apoptosis and oxidative stress injury was assessed by qRT-PCR. Compared to the control group or single factor treatment, the mRNA expression levels of *CASP6*, *CASP9*, *MAPK14*, *BAX* and *JUN* in the CDDP-WE treatment were significantly upregulated in SW620. Collectively, our findings suggest that WE may aggravate the effect of CDDP through affecting cell cycle and oxidative stress related pathways such as p53 or MAPK signaling pathway (Figure 9I).

**Figure 9.**
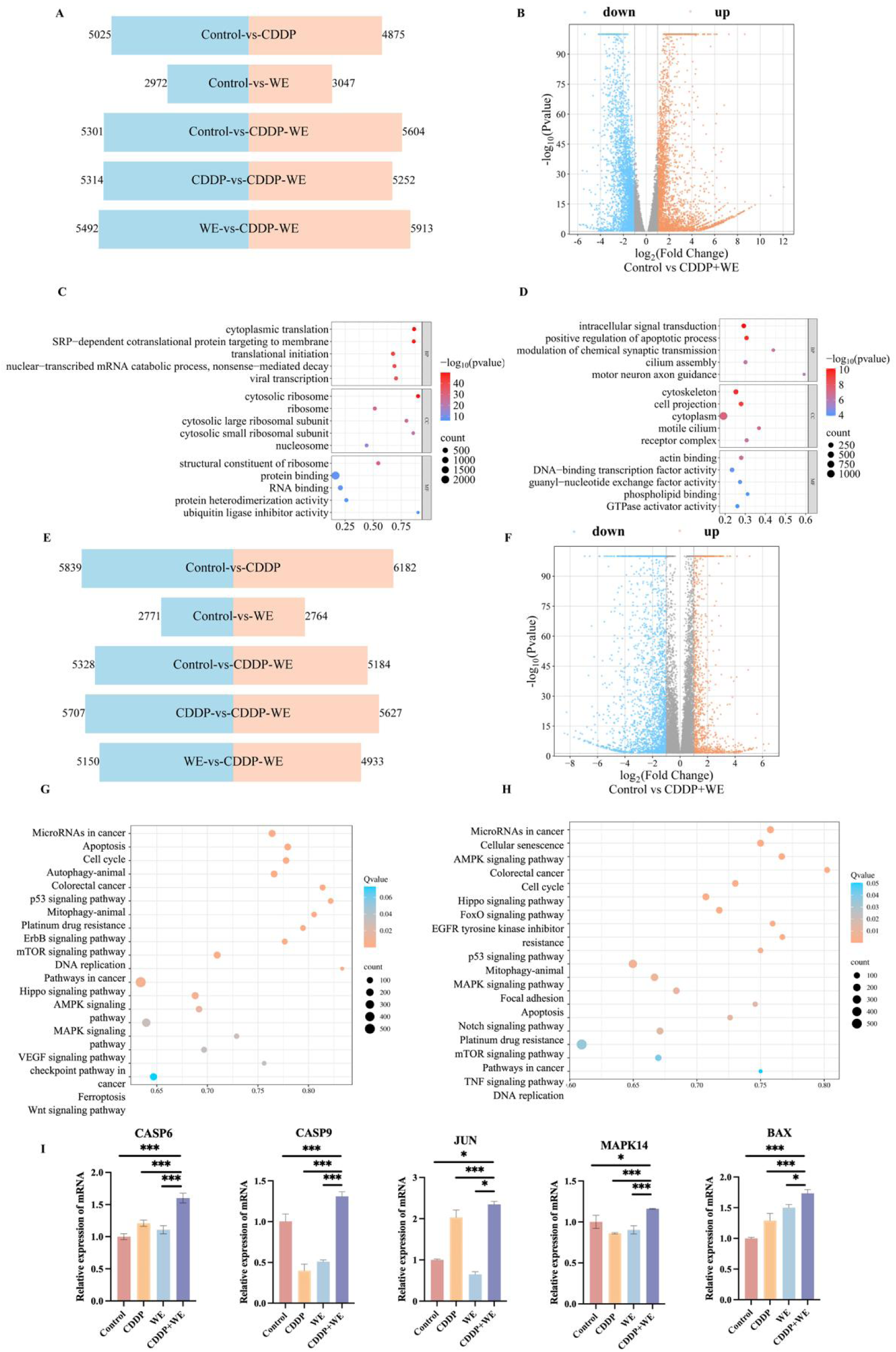
RNA sequencing (RNA-seq) analysis. (A,E) RNA-seq analysis showing the differentially modulated genes between differentially modulated genes in control vs. CDDP-WE group for both SW620 and HCT116 cell lines. (B,F) Volcano plot of DEGs between control and CDDP-WE groups, with a fold change >1 as the threshold for significant DEGs. (C-D) GO enrichment analysis of the DEGs in control vs. CDDP-WE group. (G-H) KEGG pathway enrichment analysis of the DEGs in control vs. CDDP-WE group. (I) qPCR analysis of the mRNA expression levels of selected genes in SW620. The mean ± SD is presented, n = 3. * p < 0.05, ** p < 0.01, and *** p < 0.001 vs. different group.

### DIA-based Proteomic Profiling Reveals the Impact of CDDP-WE Combination Therapy

To further validate the underlying mechanisms of the CDDP-WE combination treatment in anti-CRC synergistic effects, we investigated the impact of the CDDP-WE treatment on the global protein expression through DIA-based proteomics analysis. In total, 66,628 peptides corresponding to 8191 proteins were identified at a 1% false discovery rate (FDR) together in SW620 and HCT116 cells (Figure 10A). PCA demonstrated high reproducibility of the treatments (Figure 10B). Stringent thresholds were applied in quantitative proteomics (e.g., fold change >1.5, corresponding to log₂ FC ∼ 0.58), and we identified 133, 533, 721, 676 and 208 DEPs in the following groups: WE vs Control, CDDP vs Control, CDDP-WE vs Control, CDDP-WE vs WE, and CDDP-WE vs CDDP for the HCT116 cells. For the SW620 cells, 192, 261, 540, 363 and 269 DEPs were identified in corresponding to the above groups (Figure 10C). Notably, more DEPs were observed for the CDDP-WE treatment when compared to that by CDDP or WE alone treatment, suggesting that the combination of CDDP-WE exerts an enhanced effect in CRC cells. Clusters of Orthologous Groups (COG) analysis revealed top two pathways to be “general function prediction only” (16.60 % in SW620 and 16.05 % in HCT116) and “signal transduction mechanisms” (15.53 % in SW620 and 15.89 % in HCT116).

**Figure 10.**
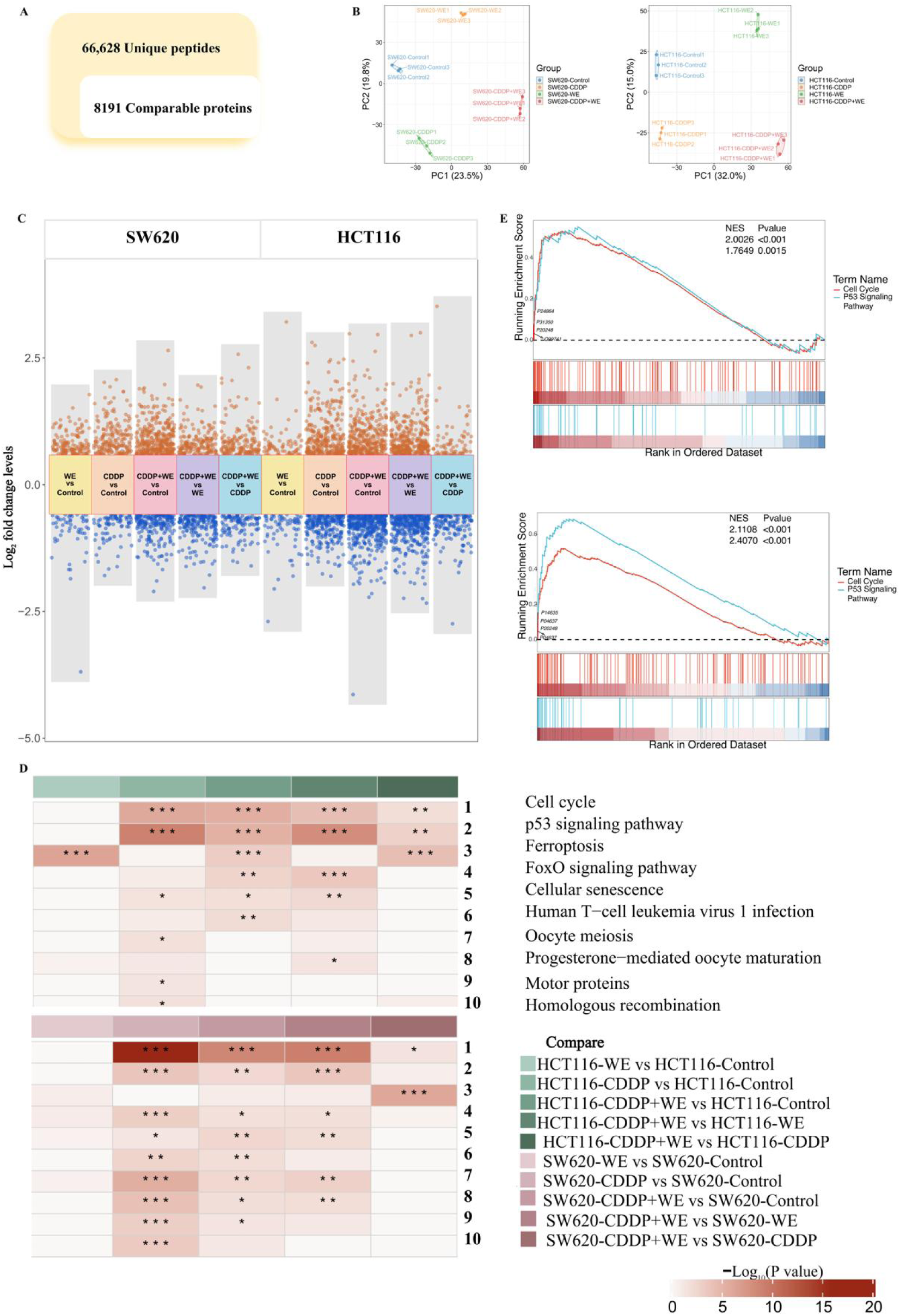
Quantitative proteomics in the SW260 and HCT116 cells treated with WE, CDDP, or combination therapy. (A) Quantitative proteome profiling identified 66,628 unique peptides, corresponding to 8191 comparable proteins. (B) Principal components analysis presentation. (C) Volcano plot of differential protein expression. (D) KEGG pathway enrichment analysis of DEGs in proteomics. (E) GESA analysis of the pathways as highlighted.

To further explore the proteomic changes related to the CDDP-WE treatment, DEPs were analyzed through GO, KEGG, and GSEA pathway enrichment analyses (Figure 10D-E). KEGG enrichment analysis confirmed the involvement of oxidative stress and other mitochondrial-related pathways such as cell cycle, P53, ferroptosis and cellular senescence pathway.

### The WE Contained a Number of Bioactive Ingredients Which Enhance the Anti-cancer Effect of CDDP

Cruciferous plants are considered as functional foods for a long time. The beneficial anticancer effects of cruciferous vegetables are generally associated with glucosinolates and their degradation products, resveratrol and its derivatives, polyunsaturated fatty acids, vitamins and minerals, which have broad potential applications in disease prevention and treatment. Studies have shown that ALA, as a promising dietary adjuvant, can enhance the efficacy of chemotherapy for colorectal cancer^25^. The potential of MSC combined with gemcitabine (GEM) chemotherapy has also been confirmed^26^. SFN and I3C have been shown to have well-known health-promoting effects^27^. Cel has shown active pharmacological activities in a variety of cancers, and clinical studies have shown its potential anticancer effect^28^. LA coordinates the dual antitumor response by promoting cancer cell apoptosis and regulating tumor microenvironment^29^. To identify active ingredients in WE with potential anti-CRC effects, we evaluated 6 prototype compounds (MSC, SFN, I3C, Cel, ALA, LA) in SW620 and HCT116 CRC cell lines. Among the candidates, SW620 and HCT116 cell viability assays revealed that all of the active ingredients exhibited significantly cytotoxicity (Figure 11). A synergistic effect was observed between CDDP and MSC in SW620 and HCT116 cells. Besides, combinations such as MSC-SFN (ZIP: 16.14), CDDP-I3C (ZIP: 15.94) and CDDP-SFN (ZIP: 4.37) exhibited synergistic inhibition only in HCT116 cells while CDDP-Cel (ZIP: 14.07) exhibited synergistic inhibition only in SW620 cells (Figure 12). This study confirms the synergism between CDDP and other anti-cancer active ingredient in selenium-enriched rapeseed. These findings suggest that SFN, MSC, SFN, I3C, Cel, ALA, LA are principal bioactive ingredients in WE for arresting cell cycle and promoting cell death.

**Figure 11.**
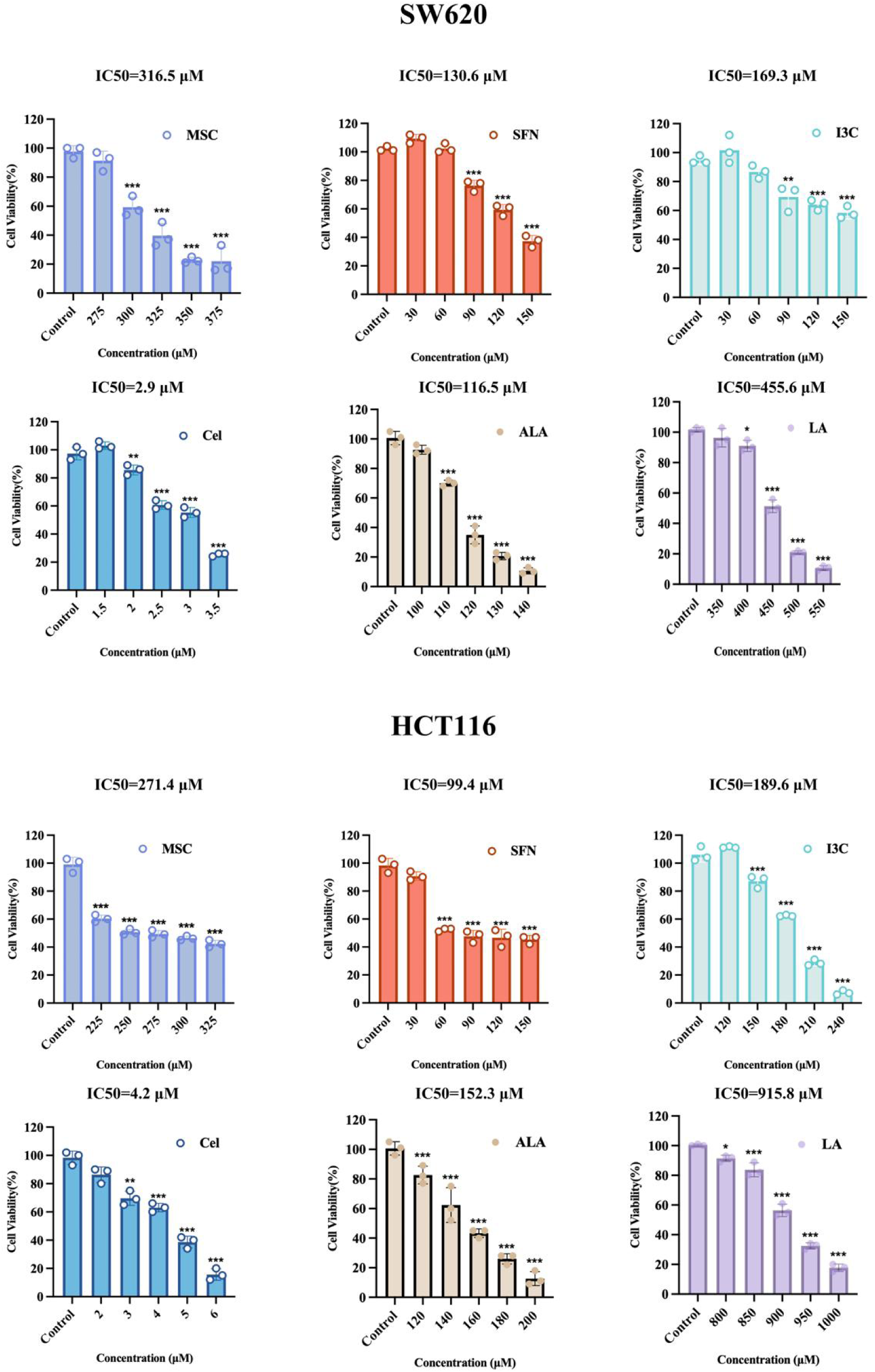
Effects of MSC, SFN, I3C, Cel, ALA and LA on SW620 and HCT116 cell viability. * p < 0.05, ** p < 0.01, and *** p < 0.001 vs. control group.

**Figure 12.**
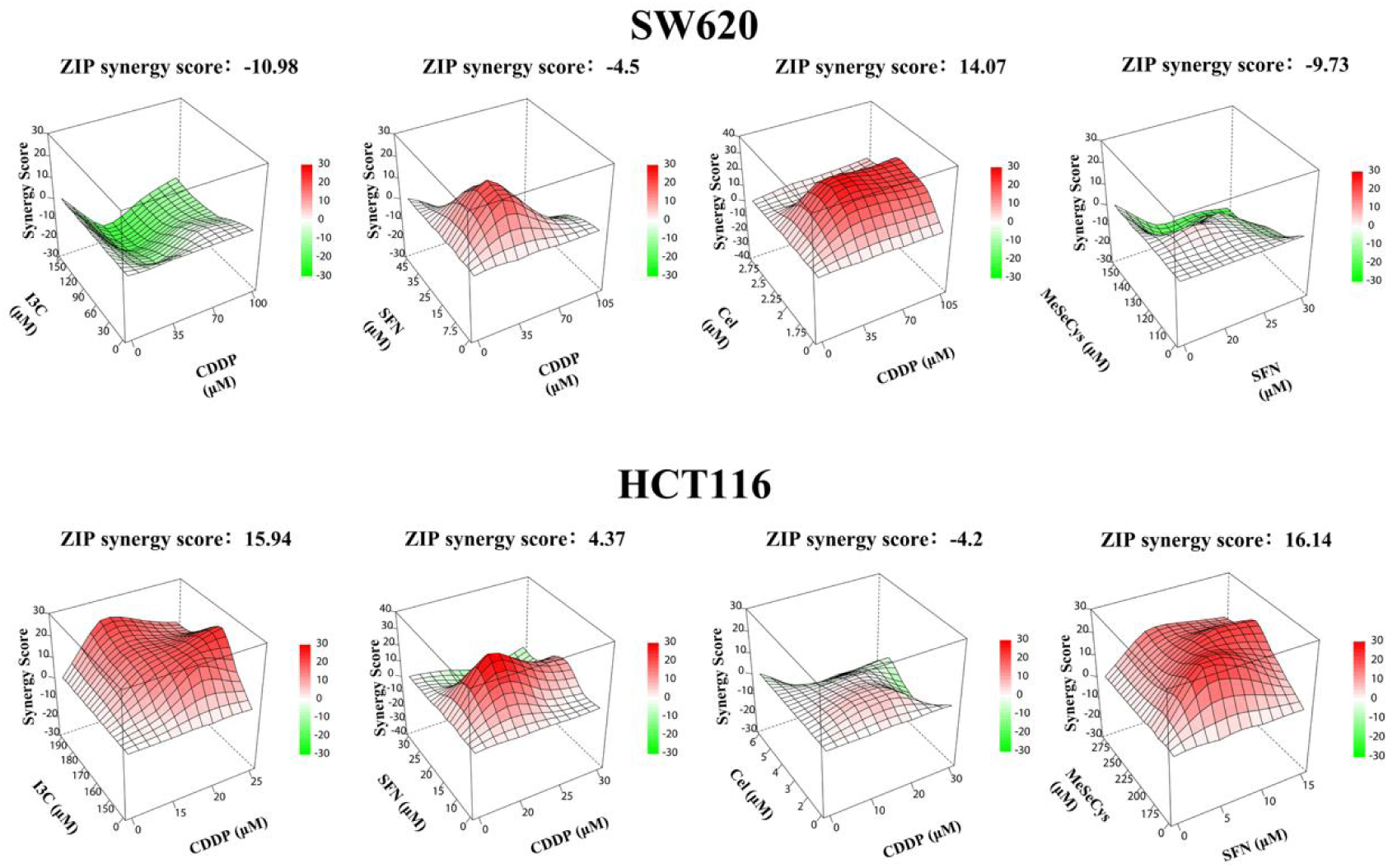
The synergy of above active ingredients contained in the selenium-enriched rapeseeds.

## DISCUSSION

In CRC treatment, traditional chemotherapy using drug such CDDP is initially effective, but later it often leads to drug resistance and disease recurrence. One way to cope with this problem is to increase the drug dosage, however, it would inevitably increase the risk of toxicity to normal cells. Therefore, a search for unharmful dietary natural products that can enhance or work synergistically with the chemotherapy drug has become a focus of extensive and ongoing research^40,41^. The huge diversities of plant metabolites are ideal resources for identifying such natural products^42^. Notably, it is often found that using the mixture of natural products often yields more efficient outcome when combined with the chemotherapy drugs. In this report, we presented strong evidence to show that the extracts from the inflorescence of the selenium-enriched oilseeds not only exerted an anticancer function alone but also enhanced the anticancer function of CDDP.

The mechanism of CDDP mainly relies on mediating oxidative stress^43^, inducing mitochondrial dysfunction^44^ and forming Pt-DNA adducts by cross-linking with DNA of CRC cells to inhibit DNA replication^45^. It also plays an important role by changing the composition of cancer microenvironment (such as recruiting macrophages, antigen presenting cells, myeloid cells and cancer-related fibroblasts) in vivo^46^. Previous studies have reported that lycopene enhances the sensitivity of oral squamous cell carcinoma to CDDP by inhibiting PI3K/Akt signaling pathway and reversing epithelial mesenchymal transition^47^. Our study revealed that the WE significantly enhanced the function of CDDP in inducing CRC cell apoptosis. In addition, the WE enhanced the function of CDDP in promoting the release of LDH, the depletion of GSH and the increase of ROS, consequently to cause an increased oxidative stress in CRC cells. Further studies showed that the WE enhanced the function of CDDP in triggering the decrease of mitochondrial membrane potential to activate the intrinsic mitochondrial apoptosis pathway.

Natural products show the potential to reverse cellular damage caused by chemotherapy drugs. For example, Au-M nanosystem elicited potent renoprotection via Nrf2-orchestrated transcriptional reprogramming in vivo^48^. Icariin (ICA) is an active flavonoid isolated from Epimedium plants. Studies have found that it can maintain the anti-cancer efficacy of CDDP while reducing its nephrotoxicity^8^. A natural food-derived ingredient-corn silk extract (CSE) can reduced the level of myocardial injury markers and provides a novel mechanistic basis aimed at supporting cardiovascular health^49^. Using LC-MS/MS systems and HPLC-ICP-MS for analysis, we found that MSC was the leading selenospecies present in WE. Subsequent cellular experiments allowed us to identify MSC, I3C, DIM, ALA, LA and SFN to be the effective ingredients that have significant inhibitory effects on CRC. Their combination with CDDP exerted a greater effect on CRC cells.

In conclusion, our studies provide evidences showing that selenium-enriched rapeseed might be a useful source for identifying natural products against CRC cells. Although the CDDP-WE combination has made promising discoveries, its practical application value still faces some challenges. Considering the long-term safety of the extracts, it is important to optimize the dose and treatment strategy, and systematic evaluation is needed. The current cell model may not fully reflect the complexity of human cancer microenvironment, and the potential side effects also need further explore in animal experiments. In addition, the immunomodulatory effect of the active ingredient combination and its combination with existing cancer treatments (such as chemotherapy, radiotherapy and targeted therapy) are worthy of further study. Clinical trials are crucial for translating these findings into practice. These limitations will also become the focus of our future research.

## NOTES

The authors declare no competing financial interest.

## ACKNOWLEDGMENTS

The authors sincerely thank Professor Jinrong Peng for valuable guidance and constructive suggestions to this study.

